# C-reactive protein and the menstrual cycle in females with sickle cell disease

**DOI:** 10.1101/2025.01.09.632177

**Authors:** Jessica Wu, Veronica Bochenek, Kandace Gollomp, Andrea H. Roe

## Abstract

There are notable sex disparities in the frequency and severity of vaso-occlusive pain episodes (VOEs) in sickle cell disease (SCD), with females experiencing more frequent and severe episodes during the reproductive years. Many individuals report a temporal link between VOEs and the menstrual cycle, suggesting that sex hormones may play a role in modulating inflammation and pain. This study explores the relationship between inflammatory markers and the menstrual cycle in females with SCD. Plasma samples from 13 female and 18 male individuals with confirmed SCD were analyzed for C-reactive protein (CRP) and other inflammatory biomarkers. In female patients, sex hormones were measured to determine the menstrual cycle phase. While no overall difference in CRP levels was observed between males and females, CRP levels fluctuated significantly across the menstrual cycle in females, with higher levels in the follicular phase compared to the luteal phase. Additionally, platelet counts were significantly elevated in females during the follicular phase compared to males. These findings suggest that cyclic inflammation may contribute to the increased frequency and severity of VOEs in females, particularly during the perimenstrual period. Further research is needed to explore whether hormonal interventions can reduce perimenstrual VOEs in SCD.

## INTRODUCTION

Sickle cell disease (SCD) is a complex disorder that exhibits sex disparities in the frequency and severity of vaso-occlusive pain episodes (VOEs). Females with SCD experience more frequent VOEs with more severe pain, leading to increased hospitalizations, particularly during reproductive years, compared to males.^1,2^ Additionally, half of females with SCD report a temporal association between VOEs and their menstrual cycle, with VOEs clustering in the perimenstrual period.^3-5^ This clinical pattern suggests that VOEs may be mediated by sex hormone cyclicity.

SCD is characterized by chronic inflammation stemming from hemolysis, endothelial dysfunction, and vaso-occlusion, with exacerbations during VOEs.^6,7^ C-reactive protein (CRP) is a key inflammatory marker that is elevated at baseline in SCD^8,9^ and rises acutely during VOEs.^10^ In comparison with a panel of hematologic and inflammatory markers, including white blood cell count, hemoglobin, lactate dehydrogenase, vascular cell adhesion molecule-1, and P-selectin, CRP emerged as the most significant biomarker correlate of VOEs requiring hospitalization.^11^ Furthermore, this robust marker of inflammation in SCD has been shown to vary across the menstrual cycle in healthy females, peaking in the follicular phase.^12-15^

Whether CRP varies by sex or fluctuates with the menstrual cycle among females with SCD has not been previously investigated. Therefore, this study aims to explore patterns of CRP and other inflammatory markers among individuals with SCD.

## METHODS

We analyzed stored plasma samples with an associated diagnostic code for SCD from the Penn Medicine BioBank repository. Using the electronic medical record, we confirmed SCD diagnosis, genotype, and reproductive age (defined as 18 to 50 years old) at time of sample collection. We excluded participants who were pregnant, hospitalized with a VOE, or being treated at an emergency department or infusion center at time of sample collection. Demographics, medications, and laboratory information (including complete blood count and complete metabolic panel) from time of sample collection were also abstracted from the electronic medical record.

We measured high sensitivity CRP in all samples and female sex hormones, including estradiol, progesterone, and luteinizing hormone, in samples from female patients. All laboratory work was performed using a Roche Cobas e411 Analyzer. Commercial ELISAs were used to quantify plasma levels of biomarkers of thrombosis (e.g. thrombin-antithrombin complexes, Abcam, von Willebrand factor, Abcam), neutrophil extracellular trap release (citrullinated histone 3, Cayman), and stress (cortisol, MP Biomedicals).

Given the absence of participant menstrual cycle information (e.g. last menstrual period and cycle length) in the electronic medical record, we predefined sex hormone cutoffs to estimate menstrual cycle phase of female participants at time of sample collection. In keeping with typical clinical laboratory definitions and our internal research in predicting menstrual cycle phase from cross-sectional hormone values, we used a progesterone level of 1.75 ng/mL to define occurrence of ovulation and thus the cutoff between follicular and luteal phases of the menstrual cycle.

We compared CRP, clinical laboratory markers, and other biomarker levels by patient sex, SCD genotype, and hydroxyurea use. Among female participant samples, we made the same comparisons between samples from the follicular and luteal phases of the menstrual cycle.

Statistical testing for comparison of medians between groups was performed using Mann-Whitney and Kruskal-Wallis tests as appropriate.

This study was approved by the Institutional Review Board of the University of Pennsylvania (protocol 851222).

## RESULTS

A total of 31 plasma samples from individuals with confirmed SCD met our inclusion criteria and were analyzed. Mean CRP concentrations in our cohort (4.45, SD 3.84 mg/ml) were consistent with levels reported in prior studies of individuals with SCD,^8,9^ and elevated compared to levels reported in the general population.^8^

No significant differences were observed in median CRP levels by SS (n=15) and SC (n=13) genotype (4.59 vs. 5.21 mg/ml, p=0.68); two patients had the S-beta thalassemia genotype and one had an unknown genotype and were therefore not included in this comparison. No difference was seen by treatment (n=6) or lack of treatment (n=25) with hydroxyurea (3.51 vs. 4.59 mg/ml, p=0.54).

Participants categorized in the follicular phase had median estradiol 64.4 pg/mL, progesterone 0.42 ng/mL, and luteinizing hormone 11.4 IU/L; those in the luteal phase had estradiol 106.9 pg/mL, progesterone 7.34 ng/mL, and luteinizing hormone 4.48 IU/L. Median CRP levels were compared by female (n=13) and male (n=18) sex, with no difference observed (3.88 vs. 4.45 mg/L, p=0.89). However, CRP did vary by menstrual cycle phase: females in the follicular phase had higher median CRP levels than those in the luteal phase (8.80 vs. 0.82 mg/L, p=0.03) (Figure 1). Although median CRP did not differ across the sexes overall, there was a trend toward higher CRP among females in the follicular phase compared with males (8.80 vs. 4.45 mg/L, p=0.59), and lower CRP among females in the luteal phase compared with males (0.82 vs. 4.45, p=0.22).

**Figure 1.**
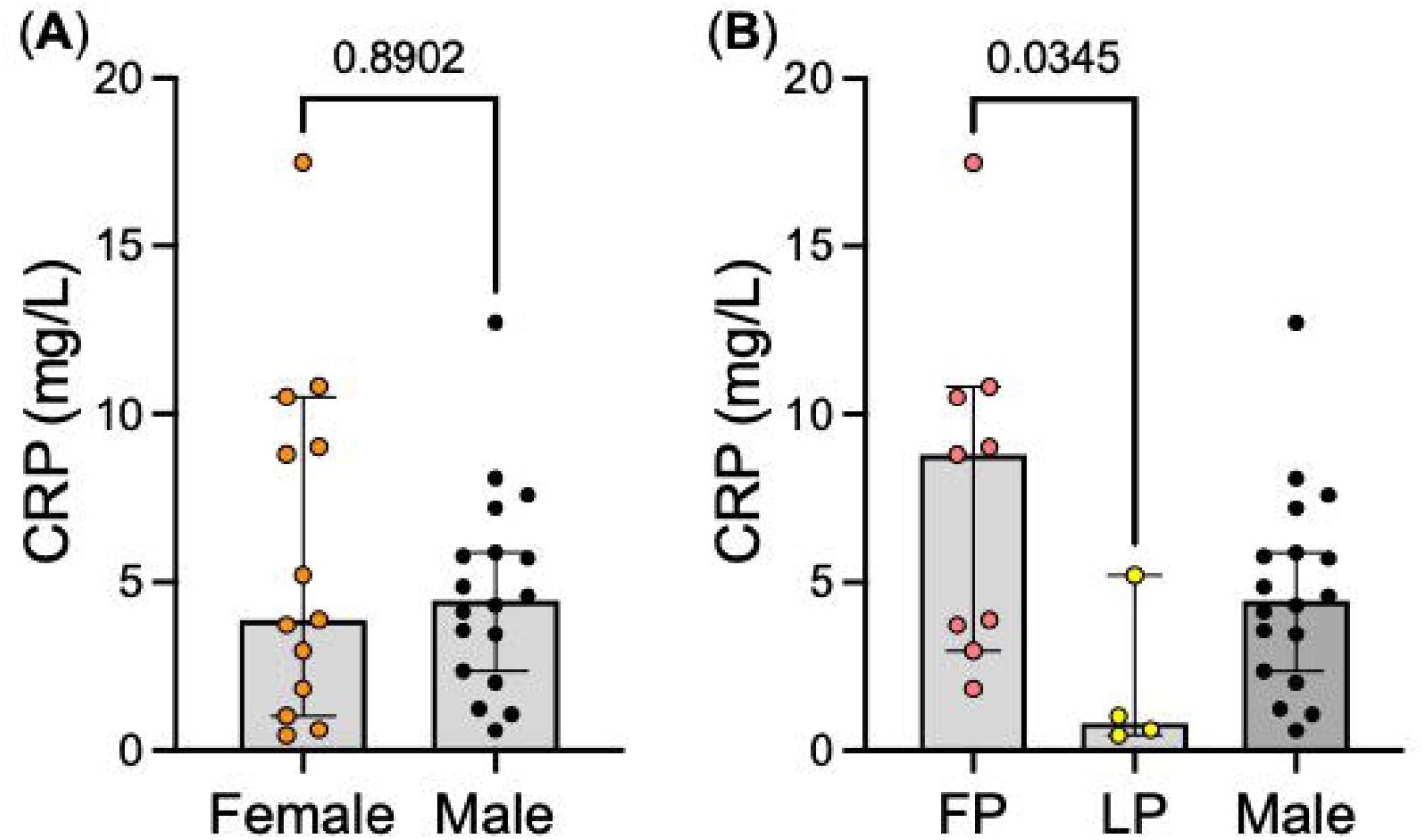
C-reactive protein (CRP) comparisons by sex and menstrual cycle phase. (A) No significant difference in CRP was detected in female (n=13) vs. male individuals (n=18) with SCD. (B) CRP was found to be significantly elevated in females in the follicular phase (FP, n=9) compared to females in the luteal phase (LP, n=4), p<0.05. No significant difference was observed between male individuals and females in the luteal phase (p=0.22) or females in the follicular phase (p= 0.59). Analysis with Kruskal-Wallis test to compare ranks between three groups.

Neutrophil and platelet counts, aspartate aminotransferase, cortisol, and thrombin-antithrombin complexes also exhibited a trend toward elevation among females in the follicular phase compared to females in the luteal phase and to males. Median platelet counts in females in the follicular phase were significantly elevated compared to platelet counts in male subjects (383k/µl vs. 219k/µl, p<0.001) (Figure 2).

**Figure 2.**
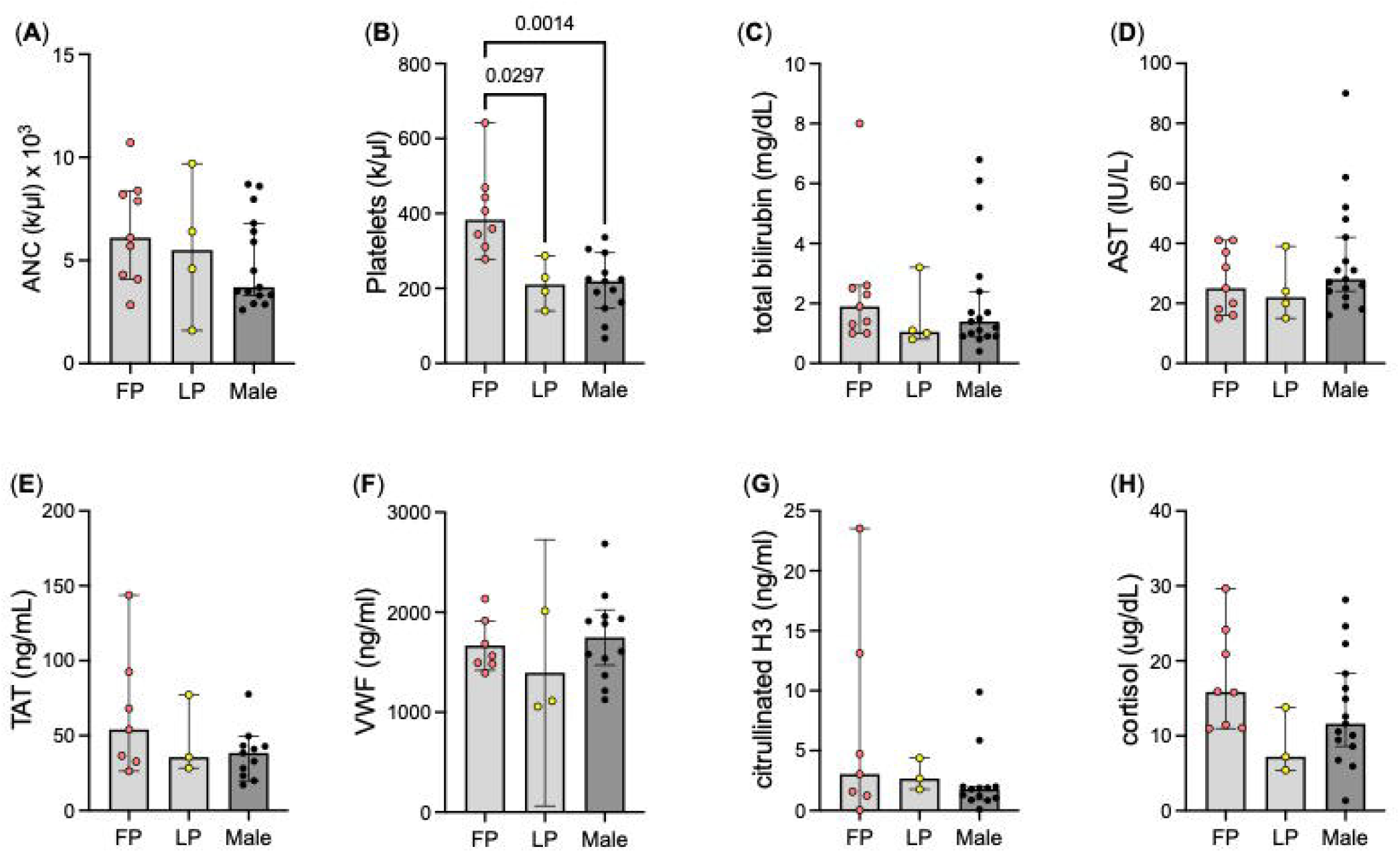
Variation in markers of cellular activation, hemolysis, thrombosis, neutrophil extracellular trap release, and stress among females and males with SCD. Cellular activation markers include (A) absolute neutrophil count, ANC, and (B) platelets. Hemolysis markers include (C) total bilirubin and (D) aspartate aminotransferase, AST. Thrombosis markers include (E) thrombin-antithrombin complexes, TAT, and (F) von Willebrand factor, VWF. Neutrophil extracellular trap release marker is (G) citrullinated histones. Stress marker is (H) cortisol. Markers were analyzed among females in the follicular phase (FP), females in the luteal phase (LP), and males with SCD. Analysis with Kruskal-Wallis test.

## DISCUSSION

CRP levels were higher in this SCD cohort compared with levels reported in the general population, consistent with previous literature.^8,9^ We observed a significant difference in CRP between females with SCD in the follicular phase compared with those in the luteal phase of the menstrual cycle, suggesting a cyclicity in inflammation that peaks in the follicular phase. This finding mirrors menstrual patterns in CRP in the general population, but CRP elevations in the follicular phase appear amplified in SCD: 8.80 in this study vs. 0.74 mg/ml in healthy females without SCD.^12^ In SCD, CRP fluctuation across the menstrual cycle may have clinical implications, as VOEs exhibit a similar temporal pattern.

This study is limited by its small sample size and cross-sectional design. Each participant contributed only one study sample, thus biomarker differences attributed to menstrual cycle phase may represent other inter-individual differences. We also lacked clinical menstrual cycle data among females; however, our use of sex hormone laboratory measurements still allowed for rigorous menstrual cycle phase definitions, while acknowledging the assumption of regular menstruation in our subjects. We also acknowledge that females with SCD have increased prevalence of diminished ovarian reserve (DOR), which is associated with low sex hormone levels; follicle stimulating hormone levels would distinguish between DOR and follicular phase of the menstrual cycle, but these were not measured. Nevertheless, individuals with DOR often have very low estradiol <20 pg/mL, which was not observed for any participant. We also lacked information regarding timing of blood draws, making it difficult to interpret differences in cortisol, which varies diurnally. Finally, because VOEs evolve through different stages and the prodromal phase is not typically associated with pain,^16^ another limitation is the inability to ensure that included subjects, while not having symptoms of an acute VOE, were at their baseline inflammatory steady state. These findings warrant prospective validation, exploration of the menstrual patterns of other markers associated with SCD pathophysiology, and correlation with clinical symptomatology.

VOEs are a crucial target for intervention among individuals with SCD, yet perimenstrual VOE pathophysiology remains poorly understood. These results suggest a cyclic pattern of inflammation across the menstrual cycle in females with SCD that may contribute to perimenstrual VOEs. Further study of perimenstrual VOEs may support therapeutic strategies, including hormonal contraceptives,^17,18^ outside of the typical armamentarium of SCD treatments.

## ACKNOWLEDGEMENTS

No sources of funding were utilized for this study.

## AUTHORSHIP CONTRIBUTIONS

Jessica Wu, MD: Involved in project creation, data collection, data analysis, and manuscript writing.

Veronica Bochenek, BA: Involved in data collection and data analysis.

Andrea H. Roe, MD, MPH: Involved in project creation, data collection, data analysis, and manuscript writing.

Kandace Gollomp, MD: Involved in project creation, data collection, data analysis, and manuscript writing.

## REFERENCES

1. Masese RV, Bulgin D, Knisely MR, et al. Sex-based differences in the manifestations and complications of sickle cell disease: Report from the Sickle Cell Disease Implementation Consortium. PLoS ONE. 2021;16(10). doi:10.1371/JOURNAL.PONE.0258638

2. Stimpson SJ, Rebele EC, Debaun MR. Common gynecological challenges in adolescents with sickle cell disease. Expert Review of Hematology. 2016;9(2):187–196. doi:10.1586/17474086.2016.1126177/SUPPL_FILE/IERR_A_1126177_SM4398.XLSX

3. Yoong WC, Tuck SM. Menstrual pattern in women with sickle cell anaemia and its association with sickling crises. J Obstet Gynaecol. 2002;22(4):399–401. doi:10.1080/01443610220141362

4. Day M, Bonnet K, Schlundt DG, DeBaun M, Sharma D. Vaso-occlusive pain and menstruation in sickle cell disease: A focus group analysis. Womens Health Rep (New Rochelle). 2020;1(1):36–46. doi:10.1089/whr.2019.0008

5. Sharma D, Day ME, Stimpson SJ, et al. Acute vaso-occlusive pain is temporally associated with the onset of menstruation in women with sickle cell disease. J Womens Health (Larchmt). 2019;28(2):162–169. doi:10.1089/jwh.2018.7147

6. Bandeira ICJ, Rocha LBS, Barbosa MC, et al. Chronic inflammatory state in sickle cell anemia patients is associated with HBB*S haplotype. Cytokine. 2014;65(2):217–221. doi:10.1016/J.CYTO.2013.10.009

7. Nader E, Romana M, Connes P. The red blood cell—Inflammation vicious circle in sickle cell disease. Front Immunol. 2020;11:454. doi:10.3389/FIMMU.2020.00454/BIBTEX

8. Schnog JB, Mac Gillavry MR, van Zanten AP, et al. Protein C and S and inflammation in sickle cell disease. Am J Hematol. 2004;76(1):26–32. doi:10.1002/ajh.20052

9. Zahran AM, Nafady A, Saad K, et al. Effect of Hydroxyurea Treatment on the Inflammatory Markers Among Children With Sickle Cell Disease. Clin Appl Thromb Hemost. 2020;26:1076029619895111. doi:10.1177/1076029619895111

10. Mohammed FA, Mahdi N, Sater MA, Al-Ola K, Almawi WY. The relation of C-reactive protein to vasoocclusive crisis in children with sickle cell disease. Blood Cells Mol Dis. 2010;45(4):293–296. doi:10.1016/j.bcmd.2010.08.003

11. Krishnan S, Setty Y, Betal SG, et al. Increased levels of the inflammatory biomarker C-reactive protein at baseline are associated with childhood sickle cell vasocclusive crises. Br J Haematol. 2010;148(5):797–804. doi:10.1111/j.1365-2141.2009.08013.x

12. Gaskins AJ, Wilchesky M, Mumford SL, et al. Endogenous reproductive hormones and C-reactive protein across the menstrual cycle: the BioCycle Study. Am J Epidemiol. 2012 Mar 1;175(5):423–31. doi:10.1093/aje/kwr343

13. Blum CA, Müller B, Huber P, et al. Low-grade inflammation and estimates of insulin resistance during the menstrual cycle in lean and overweight women. J Clin Endocrinol Metab. 2005;90(6):3230–3235. doi:10.1210/jc.2005-0231

14. Chaireti R, Lindahl TL, Byström B, Bremme K, Larsson A. Inflammatory and endothelial markers during the menstrual cycle. Scand J Clin Lab Invest. 2016;76(3):190–4. doi:10.3109/00365513.2015.1129670

15. Wander K, Brindle E, O’Connor KA. C-reactive protein across the menstrual cycle. Am J Phys Anthropol. 2008;136(2):138–146. doi:10.1002/ajpa.20785

16. Ballas, S. K., Gupta, K., & Adams-Graves, P. (2012). Sickle cell pain: A critical reappraisal. Blood, 120(18), 3647–3656. 10.1182/blood-2012-04-383430

17. De Abood MD, Castillo ZD, Guerrero F, Espino M, Austin KL. Effect of Depo-Provera or Microgynon on the painful crises of sickle cell anemia patients. Contraception. 1997;56(5):313–316. doi:10.1016/S0010-7824(97)00156-X

18. De Ceulaer K, Hayes R, Gruber C, Serjeant GR. Medroxyprogesterone acetate and homozygous sickle-cell disease. Lancet. 1982;320(8292):229–231. doi:10.1016/S0140-6736(82)90320-8

